# Acoustic Features of Emotional Expression in Preverbal Infant Vocalizations

**DOI:** 10.1101/2025.10.06.680673

**Authors:** M. Mauchand, I. Lorenzini, M. Gratier, A. Gomes-Fernandes, L. Pugin, D. Grandjean, M. Filippa

## Abstract

The musicality of human communication before the onset of words enables mutual interaction from early infancy and lays the foundation for language development. The present study investigated the development of infants’ emotional vocalizations across the first year of life, with a specific focus on their acoustic properties. Analyses revealed systematic age-related changes in acoustic features, with pronounced shifts in pitch height, spectral clarity, and formant stability occurring between 3–6 and 6–9 months. These findings mark the second half of the first year as a key milestone in the development of vocal communication. Linear discriminant analysis confirmed improved age classification beyond this stage. Emotional valence was encoded in distinct acoustic profiles, particularly involving spectral energy, pitch, vocal intensity, and resonance. Negative vocalizations, especially in younger infants, exhibited consistently higher pitch and intensity, along with stable acoustic signatures over time, suggesting their function as early, distinct affective signals. Valence-specific developmental trajectories were also identified. While formant frequency changes typically clustered around the 6-month age, an earlier shift was evident in positive vocalizations, adding nuance to general developmental trends.

Taken together, these findings challenge the notion of full emotional neutrality in preverbal vocalizations and suggest that emotional valence is partially encoded in stable acoustic parameters. Such cues may complement contextual information in supporting caregivers’ interpretation of infants’ emotional states.

This work contributes to understanding how functional flexibility emerges through the developmental tuning of prosody and further calls for research into whether adults can reliably decode emotion from vocal signals alone, shedding light on the perceptual foundations of early social communication.

## Introduction

In research on voice, affect, and emotion, the ontogeny and development of infants’ abilities to perceive voices (Bertoncini & Mehler, 1981; Blasi et al., 2011; Dehaene-Lambertz et al., 2002; Eimas et al., 1971; Paquette et al., 2018) and prosodic cues in human voices have been extensively investigated (Gervain, 2018; Gervain & Benavides-Varela, 2016). In contrast, infant’s own affective vocal productions—and the developmental trajectories of these expressive behaviors—remain comparatively underexplored (Scheiner et al., 2002; Snow, 1998; Weinberg & Tronick, 1994). Seminal work on non-segmental patterning in prelinguistic vocalisations shows that melodic organisation emerges early and is functionally oriented (D’Odorico, 1984). In D’Odorico’s longitudinal study of four Italian infants recorded twice monthly from roughly 4–5 months across a four-month window, it has been shown that the production of non-cry vocalizations in similar contexts and with recurring melodic contours reflects the infant’s emerging ability to use vocal means that are not purely physiologically driven, but functionally oriented to elicit changes in the environment - suggesting that melodic patterning may constitute the earliest form of linguistic structuring. Requests were mainly marked by rising contours, discomfort cries by falling or level contours, and calls showed intermediate patterns, suggesting that infants use melodic patterning as an early means of structuring vocalisations for communicative effect (D’Odorico, 1984).

It is now widely recognized that the vocalizations produced by infants in social contexts are an important foundation for language and communicative development (Karousou & López-Ornat, 2013; Oller et al., 2019; Papousek & Papousek, 2019; Vihman et al., 1986). These early vocalizations exhibit some important continuities with later speech and language acquisition and serve as a predictor of future spoken language abilities (LIU et al., 2023; Lyakso et al., 2014; Werwach et al., 2021). The emergence of emotional prosody in infant’s vocal utterances - despite being central to early communication - remains underexplored in acoustic research (D’Aloia et al., 2024). In particular, the acoustic analysis of the emotional valence of infant vocalizations has received relatively little attention, leaving gaps in our understanding of how infants convey and differentiate emotions through their vocal expressions across development. Emotional prosody in early vocalizations is described here as the modulation of vocal acoustic parameters to convey emotional states, intentions, and attitudes, playing a pivotal role in social communication and interpersonal interactions (Grandjean et al., 2006; Pell & Kotz, 2021). As infants gain control over their vocal tract (Kent, 2022; Locke, 2007), their vocal expression of emotion may grow richer and more complex. As early as at 3 months, functional flexibility, the capacity to produce sounds that can be used to fulfil multiple functions emerges and infants make variations on their vocalizations to express a full range of emotional valences (Oller et al., 2013). Within 6 months, melodic patterns of vocalizations complexify (Wermke et al., 2021). After 9 months, an intonational shift occurs from pre-linguistic to linguistic: intonation patterns become more guided by verbal cues and less by physiological needs and responses (Levitt, 1993). Thus, even before they can produce words, babies seem to progressively master a full intonation system and modulate their voice to express emotions and intentions. This early development of emotional prosody acts as a major interface between babies’ well-being and their caregivers: as their needs and emotions develop, so do the responses they trigger in adults (Parsons et al., 2017; Schick et al., 2022).

As the literature begins to draw clearer patterns on infants’ emotional productions, a precise assessment of the acoustic profile of these vocalizations has yet to be provided.

Previous research has acoustically described preverbal vocalisations, with studies focusing for example on segmental and suprasegmental properties of babbling, cries, and early protophones (e.g., (Iyer & Oller, 2008; Kent & Murray, 1982; Laufer & Horii, 1977). These works have provided valuable insights into developmental trajectories of pitch, formant structure, and melodic patterning. However, they have not explicitly addressed the emotional dimension of infant vocal output. To our knowledge, no systematic study has yet examined preverbal vocalisations by categorising them according to their emotional valence (positive, negative, or neutral). The present work therefore extends the scope of previous acoustic investigations by linking measurable acoustic properties to emotion-characterised vocalisations, offering new perspectives on the intersection of vocal development and early communicative functions.

Methodological inspiration can be taken from acoustic research on adult emotional speech: to describe the complex organization of acoustic parameters in emotional prosody production, researchers have worked collaboratively to identify emotional states from the voice, combining acoustic parameters from their individual studies to propose emotionally relevant baseline feature sets. One of the most popular of such sets the Geneva Minimalistic Acoustic Parameter Set - GeMAPS (Eyben et al., 2015), composed of 62 parameters (88 in its extended form) related to frequency, voice quality, spectrum, loudness, and rhythm. The GeMAPS strengths lay in its wide acoustic dimensions span, its implementation simplicity, and the importance of perceptual relevance in computing each parameter. As such, it is a great tool to robustly assess acoustic emotional features of infant vocalizations and provide meaningful interpretations.

The present study aims to investigate and deepen our understanding of the acoustic component of infants’ positive, negative, and neutral vocalisations across development during the first months of life. In particular, by examining how these emotional vocal expressions emerge and evolve over time, this research seeks to fill a gap in the literature regarding how emotional characteristics—such as valence and prosodic features—are integrated within distinct stages of vocal development. Previous phonetic research has shown that the first year is characterised by important maturational changes in vocal physiology, including increases in spectral clarity and formant frequency range (Kent, 2022; Kent & Murray, 1982; Li et al., 2021) as well as greater articulatory precision and sharper vowel definition linked to a maturing vocal tract (Robb et al., 1997; Vorperian & Kent, 2007). Building on this background, while remaining exploratory, we predicted that, in addition to known developmental patterns, infant vocalisations would also be marked by a progressive increase in emotional expressivity and specificity, indexed by emotion-specific acoustic patterns.

## Methods

### Population and recordings

The vocalisations were drawn from two corpora of infant recordings collected in naturalistic contexts. In both cases, infant–parent interactions were recorded with a digital audio recorder (Korg Sound on Sound Unlimited Track Recorder) placed near the infant, ensuring comparable audio quality and volume. Recordings were made without specific instructions for parental or infant behavior, so as to capture spontaneous exchanges. All recordings took place in the Paris region with monolingual French-speaking families, where parents exclusively addressed their infants in French.

The first corpus was composed of 1953 audio recordings lasting on average 15 minutes each. They were collected longitudinally from 24 francophone infants, aged between 1 and 5 months. From these recordings, 2266 short segments containing sound produced by infants only, with no overlap with another voice or external sound, were isolated.

The second corpus was composed of 126 audio recordings lasting approximately 30 minutes. They were collected cross-sectionally from monolingual francophone infants aged 7 to 8 months (group 1, N = 24) and 10 to 11 months (group 2, N = 18). Recordings were made using LENA audio recorders (small recording devices that are worn by the infant in a specially designed vest, cf. Canault, (Canault et al., 2016) which continuously recorded the participants’ vocal productions during 10 hours on average within a regular, at-home day. Three 30-minutes segments were retained per participant, corresponding to the three continuous segments with the highest number of vocalizations as detected automatically by LENA software analysis. All families signed written opt-out informed consent, and the procedure was approved by the Ethical Committee of the Paris Cité University.

### Segmentation of vocalizations

Recordings from both corpora were then combined and precisely segmented into single vocalizations using Praat software (Boersma & Weenink, 2018). Recordings from the first corpus constituted the basis for vocalizations in the 0-3 months and 3-6 months age ranges, while the second corpus was segmented for vocalizations in the 6-9 and 9-12 age ranges (see details below). Vocalizations of interest were defined as high-quality infants’ voiced productions, without background noise or artefacts, and lasting at least 700ms and at most 2 seconds. Non-continuous, consecutive voiced segments within this time window were considered as part of the same vocalization if they were separated by less than 300ms. During the segmentation process, experimenters aimed to capture as much emotional variability from the babies by considering all possible excerpts across the whole recording time available irrespective of their perceived salience. In case of several near-identical vocalizations (e.g., when the baby repeats the same vocalization over and over), only a few representative items were kept. In total, 693 vocalizations were segmented.

### Annotation and selection

For the segmentation, vocalizations were clustered within four age ranges to simplify analysis, based on known critical periods of vocal development: 0-3 months, 3-6 months, 6-9 months, 9-12 months. Most selected utterances were from babies in the center of a range (1-2 months, 4-5 months, 7-8 months, and 10-11 months). During the segmentation, the following information was extracted from the vocalizations: age and age range (0-3 months, 3-6 months, 6-9 months, 9-12 months), sex of the baby, duration of the vocalization, and estimated valence of the vocalization (Positive, Neutral, or Negative). This estimation was based on the qualitative evaluation of the vocalization within its auditory context (e.g. parent’s speech) by the experimenters. Vocalizations where qualified as negative when preceded by a negative event (e.g. parent scold, loud noise) or prompting a comfort response from the parent, or preceding/following a crying phase; positive when elicited during play or accompanied by positive speech from the parent; neutral when elicited in the absence of a parent or stimulation. This categorization was performed by two trained master’s students supervised by an expert researcher who reviewed and validated each annotation. The students did not have prior knowledge of the specific predictions of the study. In cases of disagreement, the vocalization was discarded. Details of each vocalizations are available in the Supplementary Materials.

In a second phase, the segmented vocalizations were more narrowly selected to form a harmonized database. The selection process followed the criteria below: 90 vocalizations per age range with 30 positive, 30 neutral, and 30 negative vocalizations in each range, balanced sex across age ranges, and similar duration statistics across age range (M = 1.2s, SD = .3s). To account for volume variability across the recordings, all the selected stimuli were normalized to a mean sound pressure of 70dB using Praat software (Boersma & Weenink, 2018). In total, 360 vocalizations (4 age ranges x 3 valences x 90 stimuli) were selected. Examples of the selected vocalizations are displayed as spectrograms in Figure 1.

**Figure 1.**
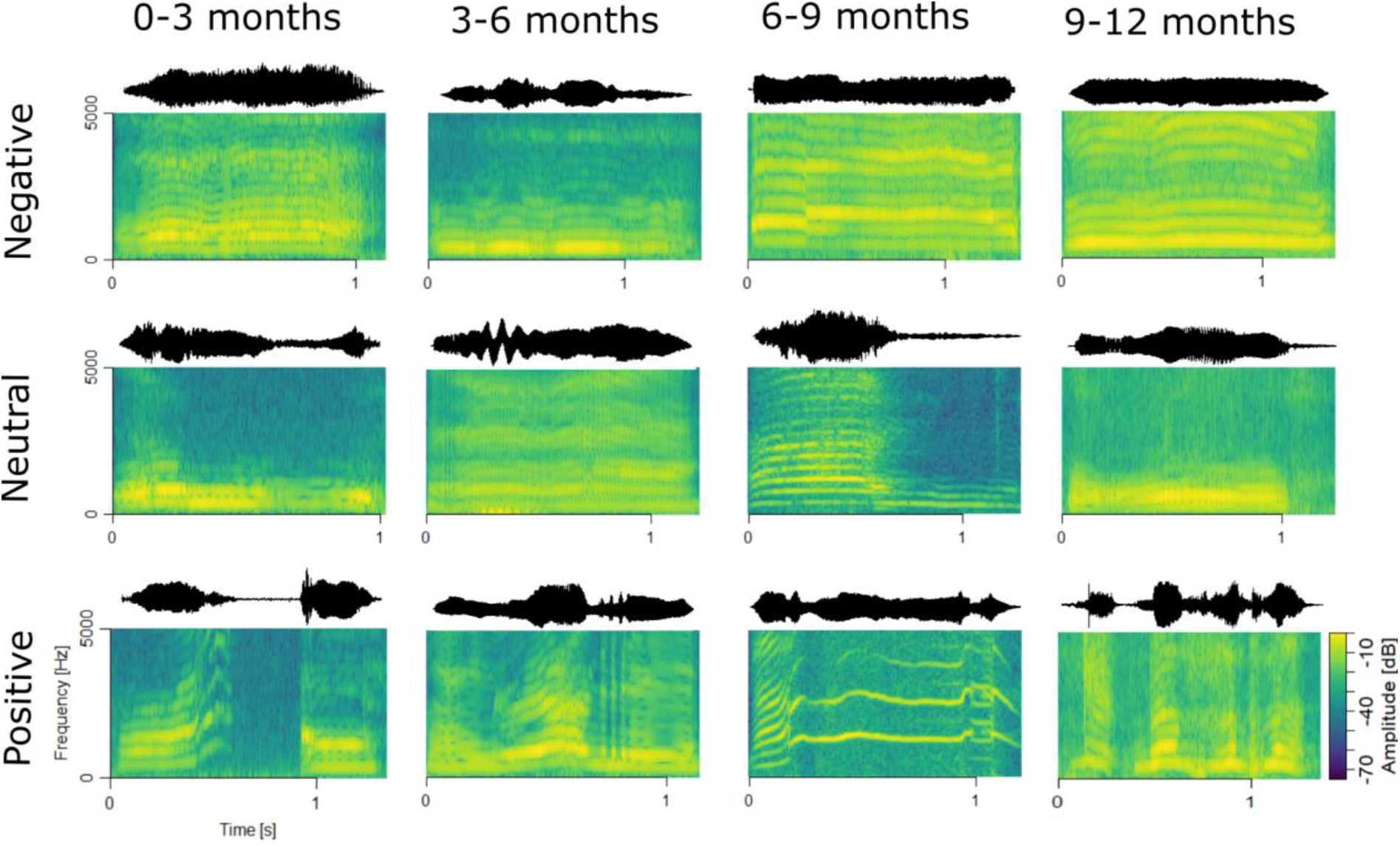
Spectrograms of selected vocalizations in each condition. The sound waveform of each vocalization is also included above the spectrogram.

### Acoustic Analyses

Acoustic parameters were extracted using the extended Geneva Minimalistic Acoustic Parameter Set (eGeMAPS, Eyben et al., 2016) in openSMILE software (Eyben et al., 2010). The eGeMAPS is a set of 88 acoustic parameters related to the frequency, loudness, spectrum, and rhythm, which are relevant to emotional speech analysis. The parameters were extracted dynamically across sound samples, and then passed through mathematical functions (e.g., mean, standard deviation, maximum) applied to the whole duration of the vocalization. The complete dataset of eGeMAPS values extracted for each vocalization is available in the Supplementary Materials.

To reduce these 88 parameters to a restricted number of interpretable values, a Principal Component Analysis (PCA) with varimax rotation was performed on the database, resulting in 5 Rotated Components (RC). The PCA scores of each vocalization was then entered into the statistical models described below.

To describe the developing acoustic profile of emotional vocalizations, the scores of each vocalization for each RC were entered into separate two-way ANOVAs, with Age Range (0-3, 3-6, 6-9, 9-12 months) and Valence (Negative, Neutral, Positive) as interacting fixed factors.

To assess the ability of the acoustic features to predict the age and emotion of a vocalizing baby, the scores of each vocalization for all RCs were entered into a Linear Discriminant Analysis. In this analysis, Age Range and Valence were collapsed together to form 12 independent Conditions of a single dependent variable (0-3 Negative, 0-3 Neutral, 0-3 Positive, 3-6 Negative, 3-6 Neutral, etc.), to be classified by a linear combination of the 5 predictors. Model predictions were then tested using leave-one-out cross-validation.

## Results

The principal components analysis of the 88 eGeMAPS features extracted from all vocalizations revealed five rotated components, each related to a different family of acoustic features: spectral energy and pitch-related features (RC1), vocal intensity and resonance (RC2), formant structure and variability (RC3), spectral bandwidth and stability (RC4), voice quality and harmonicity (RC5). Together, these components explained 43% of the acoustic variance across vocalizations. Table 1 summarizes the components and their main parameters; the complete PCA results can be found in Table S1 of the supplementary material.

**Table 1.** Summary of Principal Components (RCs) with Explained Variance, Key Loadings, and Interpretations of Acoustic Features.

| RC | Explained Variance (%) | Top Positive Loadings | Top Negative Loadings | Description | Interpretation |
| --- | --- | --- | --- | --- | --- |
| RC1 | 36.26 | Alpha Ratio: 0.89<br>Spectral slope: 0.80<br>F0 Mean: 0.784<br>F0 50<sup>th</sup> percentile: 0.73<br>F0 80<sup>th</sup> percentile: 0.71 | MFCC1 (voiced): -0.85<br>MFCC1 (all): -0.84<br>Hammarberg Index: -0.78<br>F1 bandwidth: -0.70<br>Harmonic difference H1-A3: -0.66 | Spectral energy and pitch-related features. | Captures spectral energy distribution and fundamental frequency characteristics. Positive loadings relate to spectral slope and pitch measures while negative loadings primarily involve spectral envelope (MFCC) features. |
| RC2 | 22.18 | Loudness 20<sup>th</sup> percentile: 0.76<br>F1 amplitude: 0.75<br>F2 amplitude: 0.75<br>F3 amplitude: 0.74<br>Loudness 50<sup>th</sup> percentile: 0.63 | Loudness sd: -0.81<br>Spectral flux sd: -0.754<br>Unvoiced segment length M: -0.61<br>Unvoiced segment length sd: -0.61 | Vocal intensity and resonance. | Represents vocal intensity and resonance characteristics. Positive loadings relate to loudness and formant amplitudes, whereas negative loadings capture variability in loudness and spectral features. |
| RC3 | 11.98 | F1 frequency: 0.77<br>F3 frequency: 0.74<br>F2 frequency: 0.65 | F2 frequency sd: -0.72<br>F1 frequency sd: -0.64 | Formant structure and variability. | Captures formant structure and frequency characteristics. Positive loadings represent mean formant frequencies; negative loadings reflect their variability. |
| RC4 | 10.3 | F2 bandwidth: 0.69<br>F3 bandwidth: 0.67<br>MFCC4 (voiced): 0.64<br>MFCC4 (all): 0.61 | F2 bandwidth sd: -0.67 | Spectral bandwidth and stability. | Represents spectral bandwidth characteristics. Positive loadings relate to mean bandwidth measures and specific spectral features (MFCC4), while negative loadings capture bandwidth variability. |
| RC5 | 19.28 | Jitter: 0.74<br>Shimmer: 0.74<br>Spectral flux (voiced): 0.71<br>Spectral flux (all): 0.70<br>F0 sd: 0.67 | HNR: -0.83 | Voice quality and harmonicity. | Captures voice quality measures. Positive loadings relate to perturbation (jitter, shimmer) and spectral variability, while the strong negative loading (HNR) indicates decreased harmonicity. |

### RC1, spectral energy and pitch

RC1 relates to changes in vocal definition, with respect to pitch, spectral slope, and spectral envelope. It appears to capture the interplay between the clarity and energy distribution in the vocal signal (Scherer, 2003; Zwicker & Fastl, 2013). Higher RC1 values correspond to vocalizations with clearer and higher pitch information, characterized by greater harmonic energy and stability in fundamental frequency (F0) measures; they also indicate a less pronounced spectral envelope, suggesting reduced dominance of noise-like characteristics and improved harmonic clarity (Tamarit et al., 2008).

### RC2, Loudness and resonance

RC2 reflects vocal intensity, resonance, and the degree of vocal variability and continuity. High positive loadings on these measures indicate that elevated RC2 scores are associated with vocalizations characterized by consistently strong intensity and prominent resonances. High formant amplitude relative to the fundamental frequency (F0) conveys distinct vowel-like qualities, indicating resonance in the vocal tract’s filtering characteristics (Ito et al., 2010). In terms of variability, low acoustic variability is a key feature of high RC2 scores. Negative loadings imply that as RC2 increases, vocalizations exhibit steadier loudness and a more stable spectral profile (Scherer et al., 2003).

### RC3, Formant structure and variability

This component reflects the height, clarity and stability of formant frequencies, with higher RC3 scores corresponding to higher mean frequencies of the first three formants, with reduced variability on these frequencies. Formant frequencies shape human vocalizations as indicators of vowel sounds, modulated in particular by vocal tract shape and size, as well as vocal tension (Gelfer & Mikos, 2005).

### RC4, Spectral bandwidth

This component reflects a composite measure of the spectral envelope’s breadth and spectral shape, integrating characteristics of formant bandwidth and MFCC-derived spectral features. High positive loadings for the average bandwidths of F2 and F3 suggest that broader bandwidths in these formants contribute to higher component scores, indicating less sharply defined resonances in these frequency regions, accompanied by more homogenized spectral captured by MFCC4 loadings.

### RC5, Voice quality and instability

This component captures aspects of vocal stability and periodicity in contrast to variability and noise within the acoustic signal. Positive loadings on measures such as jitter, shimmer, and spectral flux indicate that higher scores on this component are associated with greater variability and irregularity in the voice. Conversely, the strong negative loading on Harmonics-to-Noise Ratio (HNR) suggests that lower component scores correspond to a more periodic, harmonic, and stable vocal signal (Patel et al., 2010).

### ANOVAs of PCA scores

The acoustic profile of vocalizations from PCA scores is displayed in Figure 2. A summary of the scores is also available in Table S2 of the Supplementary Materials.

**Figure 2.**
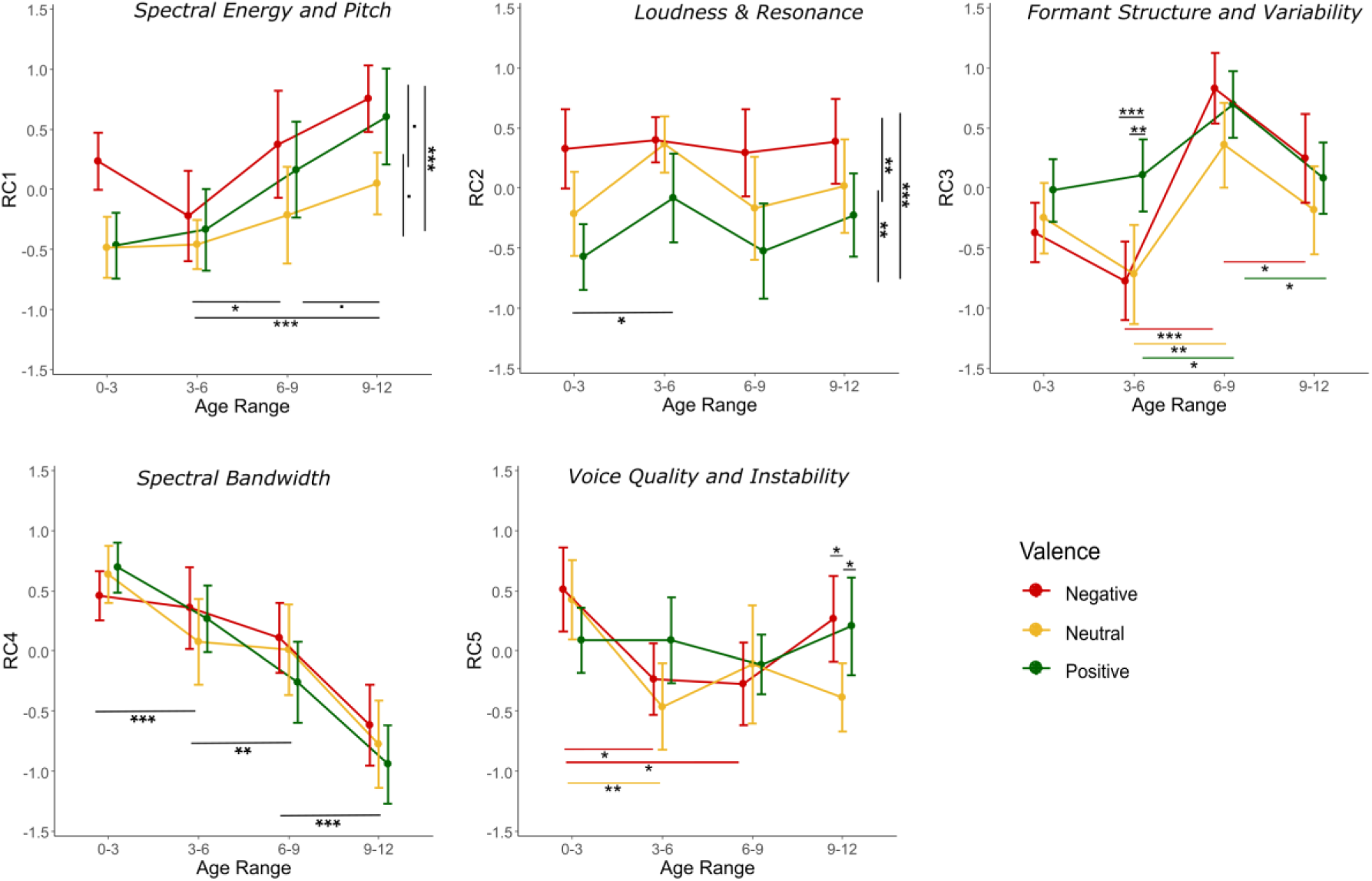
Average acoustic PCA scores for each valence type at each age range. Each panel represents the scores for one of 5 components from the acoustic PCA. The title of the panel refers to the perceptual correspondence of the component. Error bars represent 95% confidence intervals. Significance: * p < .05, ** p < .01, *** p < .001. The colour of the significance bars corresponds to the valence being tested (black = test across valences).

For RC1 (spectral energy and pitch), the model showed significant, medium main effects of Valence (F(2, 348) = 14.15, p < .001, η2 = .06) and Age Range (F(3, 348) = 11.06, p < .001 η2 = .11), with no interaction. Post-hoc tests with Holm correction revealed that Negative vocalizations exhibited significantly higher RC1 scores than Neutral ones (t(229) = 4.72, p < .001), while positive vocalizations scores were situated in-between, marginally lower than for Negative (t(237) = −2.18, p = .060) and higher than Neutral (t(223) = 2.16, p = .060). In terms of age, scores were similar for the 0-3 months and 3-6 months ranges (t(176) = .43, p = .43), with a significant increase in the 6–9-month range (t(165) = 2.88, p = .018) which continued marginally in the 9-12 months range (t(169) = 2.30, p = .063).

For RC2 (loudness and resonance), the model showed a significant effect of Valence (F(2, 348) = 16.23, p < .001, η2 = .09) and a significant, but smaller, effect of Age Range (F(3, 348) = 3.19, p < .024, η2 = .03). Post-hoc tests revealed that Negative vocalizations exhibited higher RC2 scores than both Neutral (t(233) = 2.88, p = .009) and Positive (t(234) = 5.84, p < .001) ones, and Neutral vocalizations had higher scores than Positive ones (t(238) = 2.71, p = .009). In terms of age, the only effect was significantly higher scores at 3-6 months compared to 0-3 months (t(172) = 2.92, p = .024).

For RC3 (formant structure and variability), the model showed a large effect of Age Range (F(3, 348) = 24.64, p < .001, η2 = .18) and a small effect of Valence (F(2, 348) = 6.53, p = .002, η2 = .04), with a significant interaction (F(6, 348) = 2.62, p = .017 η2 = .04). Post-hoc tests revealed that no significant difference was found between the three Valence conditions at 0-3 months (ps > .18). At 3-6 months, Positive vocalizations showed higher RC3 scores than both Negative (t(58) = 3.91, p = .001) and Neutral ones (t(53) = 3.18, p = .005). There was a significant increase at 6-9 months for all conditions, Negative (t(57) = 7.20, p < .001), Neutral (t(57) = 3.89, p = .002), and Positive (t(58) = 2.81, p = .027), and differences between the three conditions were reduced to non-significance at that age (ps > .15). Finally, there was a decrease at 9-12 months, only significant for Negative (t(55) = 2.43, p = .037) and Positive (t(58) = 2.96, p = .022) vocalizations, but no significant difference was found between Valence conditions (ps > .33).

For RC4 (spectral bandwidth), the model only showed a large effect of Age Range (F(3, 348) = 40.60, p < .001, η2 = .26). Scores across all valences showed a decrease from 0-3 to 3-6 months (t(156) = −3.15, p = .004), to 6-9 months (t(178) = −2.02, p = .045), and finally to 9-12 months (t(178) = −5.16, p < .001).

For RC5 (voice quality and instability), the model showed a small effect of Age Range (F(3, 348) = 5.91, p = .001, η2 = .05), no effect of Valence (F(2, 348) = 1.73, p = .178, η2 = .01), but a significant interaction (F(6, 348) = 2.25, p < .038, η2 = .04). Post-hoc tests showed a decrease in RC5 scores from 0-3 to 3-6 months for Negative (t(56) = −3.16, p = .015) and Neutral (t(58) = −3.57, p = .004) vocalizations; this effect continued at age 6-9 for Negative vocalizations only (t(58) = −3.14, p = .015). At age 9-12, Neutral vocalizations were found to have lower scores than both Positive (t(52) = −2.34, p = .046) and Negative (t(55) = −2.81, p = .021) ones.

### LDA classification from PCA scores

The Linear Discriminant Analysis aimed to classify the 12 Conditions, each with a prior probability of 8.33%, via linear combinations of the 5 RCs. The model extracted five linear discriminants (LD); the coefficients of each RC in the LDs is displayed in Table 2. Testing the LDA model using leave-one-out cross-validation resulted in an accuracy of 21.94%, significantly greater than the No Information Rate of 8.33% (p < .001, κ = .15). The confusion matrix shown in Figure 3 reveals that despite each Condition being entered independently, the model frequently predicted either the correct age, or the correct emotion. In most cases, confusion occurred within one Age Range, going to or from Neutral valence. A clear age effect suggests increased classification accuracy after 6 months: for both 6-9 months and 9-12 months age ranges, the most often predicted Valence was the correct one. Meanwhile, 0-3 and 3-6 Age Ranges showed more misclassifications, with especially low accuracy in the 3-6 range. Finally, Negative vocalizations were often misclassified as Negative in the wrong Age Range, suggesting a more discriminable, age-invariant acoustic profile for negative vocalizations.

**Figure 3.**
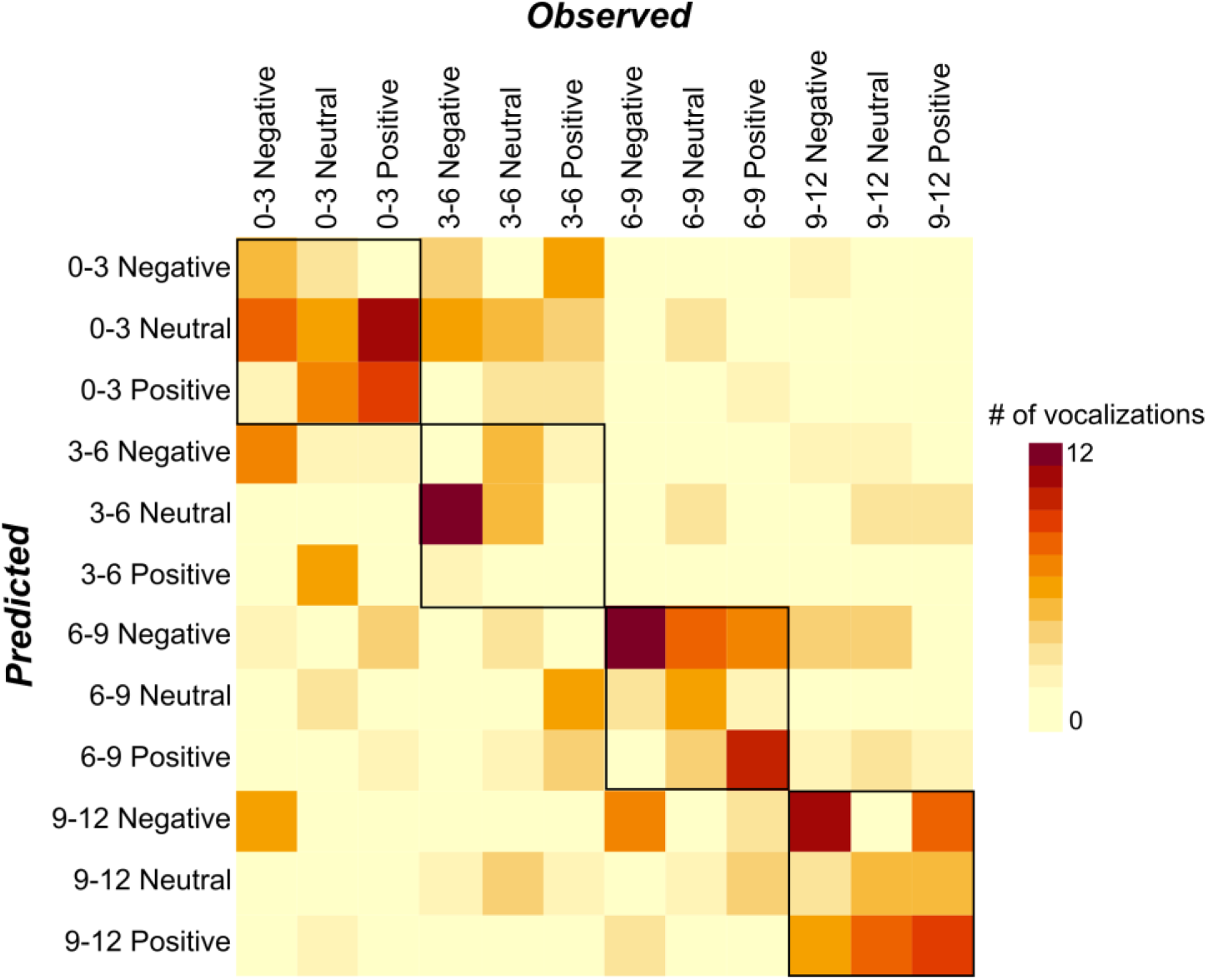
Confusion matrix for the predictions of the Linear Discriminant Analysis of PCA scores. Each square represents the number of items predicted by the model as the condition on the y-axis that were observed as the condition in the x-axis (darker hues corresponding to greater numbers). Fully correct classifications are visible on the diagonal, and Age-correct classifications are seen inside the black boxes.

**Table 2.** Summary of the Linear Discriminant Analysis.

| Linear discriminants | LD1 | LD2 | LD3 | LD4 | LD5 |
| --- | --- | --- | --- | --- | --- |
| Proportion of trace | 59.62% | 21.83% | 10.58% | 7.23% | 0.74% |
| *PCA components coefficients* |  |  |  |  |  |
| RC1 | -0.72 | 0.15 | 0.53 | 0.04 | -0.65 |
| RC2 | 0.01 | 0.59 | 0.40 | 0.64 | 0.43 |
| RC3 | -0.61 | -0.81 | 0.03 | 0.43 | 0.28 |
| RC4 | 0.91 | -0.43 | 0.40 | 0.33 | -0.34 |
| RC5 | 0.03 | -0.17 | 0.71 | -0.60 | 0.43 |

## Discussion

The present study identified significant effects of both developmental age and emotional valence across five principal acoustic components in infant’s preverbal vocalizations during the first year of life: spectral energy and pitch, vocal intensity and resonance, formant structure, spectral bandwidth, and voice quality. These effects reveal important developmental changes in infants’ emotional expression through the first year of life, prior to the acquisition of language.

### Age effect on infants’ vocal development

The longitudinal examination of vocalizations revealed critical acoustic changes related to infants’ development. The starkest changes appeared between the 3-6- and 6-9 months range, with a clear shift in spectral energy and pitch-related features, and in formant structure and their variability, across emotional conditions. This shift indicates major increases in the height, clarity, and stability of infants’ pitch, spectral structure, and formant frequencies. Such increases may be related to growth-related changes in vocal maturity and control, consistent with previous studies indicating an increases in spectral clarity and formant frequency range in this period (Kent & Murray, 1982; Li et al., 2021). Interestingly, the pronounced acoustic modifications taht were observed between 3–6 and 6–9 months may indeed be interpreted in relation to the developmental milestone marking the transition from predominantly non-intentional to increasingly intentional communication. This age-related increase may in fact reflect both a rise in arousal in dyadic interactions and the emergence of new expressive abilities (Wass et al., 2022). Around six months infants become increasingly capable of conveying signals and intentions to social partners with greater intensity and precision (Camaioni, 2017), developing the ability to co-ordinate attention toward an object or event and a social partner. This is an important step towards the achievement of an intentional form of communication.

Moreover, the present results align with findings by D’Aloia (D’Aloia et al., 2024), who reported a gradual increase in F₀ mean of approximately 3.2 Hz per month across pre-lexical vocalisations from 4 to 16 months. This gradual increase contrasts with studies that found stable or fluctuating values within the first year (Iyer & Oller, 2008; Laufer & Horii, 1977), but is partially in line with earlier observations of rising F0 during infancy (Fairbanks, 1942; Murry et al., 1983). Taken together, these findings suggest that while some investigations describe F0 as relatively stable across early development, others—especially those employing dense longitudinal sampling—point to a subtle but measurable upward trend. This divergence may reflect methodological differences in sampling frequency, inclusion criteria for vocal types, and the treatment of individual variability, which D’Aloia et al. (2024) highlighted as a central factor in explaining inconsistencies across studies.

Our findings further support the notion that around 6 months of age marks a major developmental milestone in the trajectory of human vocal expression (Kuhl et al., 2008; Nathani et al., 2006). This critical period was followed by feature-specific adjustments in the 9-12 months range, with a further marginal increase in pitch and spectral clarity compensated by a small drop in formant frequency, potentially reflecting the refining of vocal expression as infants approach speech readiness (Kuhl et al., 2008). Interestingly, these characteristics would then remain relatively stable until 24 months of age, when new developmental changes would lead to drops in formant frequencies (Gilbert et al., 1997). This consolidates the 6–12-month prelinguistic window as a critical period in vocal development.

The Linear Discriminant Analysis confirmed that acoustic profiles could partially predict age, with classification accuracy improving markedly after 6 months. The 6-month milestone was further exemplified in the linear discriminant analysis, as vocalizations from infants older than 6 months were largely classified accurately, and almost always classified to their correct age range. Meanwhile, younger infants produced less age-related discriminable vocalizations, resulting in a lot of confusion and classification porosity between 0-3- and 3-6-months ranges. Hence, the acoustic changes described at 6 months may not only reflect a general developmental trend in vocal growth but point to new emotional and interactive abilities of infants, who then begin to use more prototypical expressions through their second semester of life (Gaffan et al., 2010).

A few other developmental changes do seem to occur prior to 6 months, such as a small increase in loudness and resonance between 3 and 6 months. This localized shift may reflect an increase in vocal power due to muscular tone (Hsu et al., 2000), later compensated for by the new vocal control skills acquired at 6-months. Infant development was also marked by a steady decrease in formant bandwidth and spectral variability. These changes, indicating sharper vowel definition, are associated with greater articulatory precision and a more mature vocal tract configuration (Robb et al., 1997; Vorperian & Kent, 2007). The consistent decline in spectral bandwidth reflects a form of resonance tuning and vocal control in a more gradual manner than the changes occurring during the 6-months milestone. This developmental refinement, most evident after 6 months, is in line with the notion that the ability to imitate prosodic contours, such as those of the maternal voice (Gratier & Devouche, 2011).

### Emotion-related acoustic patterns emerge across development

The emotional valence of infant vocalizations was reflected in distinct acoustic profiles across multiple parameters, underscoring infants’ communicative abilities even before the emergence of language. Clear emotion-related patterns were revealed regarding acoustic components spectral energy and pitch-related features, and vocal intensity and resonance.

For spectral energy and pitch-related features, negative vocalizations exhibited the highest values, characterized by strong pitch-related features, a steeper spectral slope, and a more articulated signal, suggesting that, irrespectively of age, negative vocalizations are performed with great vocal precision and a typical high-pitch signature, thus potentially facilitating their discernment (Eyben et al., 2010). Positive vocalizations displayed intermediate spectral energy and pitch-related features values, balancing features of both negative and neutral vocalizations, while neutral vocalizations had the lowest scores, marked by weaker pitch definition and flatter spectral slopes. This pattern may reflect a relationship between spectral energy and pitch-related features and emotional arousal, with more emotionally expressive vocalizations exhibiting higher, clearer, and more harmonic pitch signals, especially in the context of parent-infant communication (Trainor et al., 1997; Wang et al., 2018).

Similarly, the loudness and resonance of vocal expressions showed the highest values for negative vocalizations, revealing their consistently high vocal intensity, clearer formant resonances, and reduced variability in loudness and spectral flux. This suggests that negative emotional expressions are acoustically louder, resonant, and sustained, contrasting sharply with positive vocalizations, while neutral ones exhibited more moderate, in-between scores on this parameter. Complementing the arousal pattern interpretation, loudness and resonance parameters may here relate to a valence spectrum, going from negative, loud and resonant vocalizations to positive, softer ones. Most negative vocalizations, being distress or discomfort signals, may express this distress through increased pressure in the vocal tract to alert their caregiver, even from afar (Lingle et al., 2012; Soltis, 2004). Meanwhile, positive vocalizations are typically produced in the close presence of a parent and in a pleasant context appropriate for less loud, softer productions (Kamiloğlu et al., 2025).

This recurrence of defined acoustic patterns across different emotional contents was confirmed by the linear discriminant analysis, which yielded relatively accurate classification of emotional valence. Notably, even when vocalizations were misclassified in terms of age, they were often still correctly identified with respect to their emotional valence, albeit within a different age group. This suggests the presence of an age-invariant acoustic signature for emotion, meaning that certain emotional cues in vocalizations remain perceptually stable across developmental stages.

This was particularly true for negative vocalizations, which stood out due to their distinct spectral energy, elevated pitch, increased loudness, and strong resonance - features that consistently placed them at the extreme ends of the acoustic spectrum, thus enhancing their discriminability. Interestingly, these same acoustic patterns are also characteristic of emotional expressions in adult speech, where high arousal and negative valence are similarly encoded through comparable acoustic cues (Scherer et al., 2017). As such, even prior to language acquisition, infants begin to develop abilities to communicate acoustically distinct emotional signals with illocutionary force, beyond extreme acts such as crying.

Taken together, these findings question the assumption that non-cry preverbal vocalisations are emotionally neutral, as they indicate that even early in development such vocalisations convey systematic acoustic markers of emotional valence.

### Developmental Trajectories of Emotional Valence in Infant Vocalizations

After first addressing age-related changes in early vocal development and then highlighting the emergence of distinct acoustic patterns associated with emotional valence, we examined how these emotional expressions themselves evolve across developmental time. While our findings are consistent with previous research on general age-related trends in infant vocal development (Kuhl et al., 2008; Nathani et al., 2006), they also extend this work by adding an emotional dimension - specifically, by examining how the emotional valence of vocalizations evolves throughout early development. Save for a few studies, most acoustic research on infant vocal affect is limited to infant cry (Fort & Manfredi, 1998). Our data, expanding on non-cry, emotionally diverse vocalizations, highlights several valence-specific trajectories which complexify the assessment of infant vocal development.

The analysis of formant structure and variability suggests that the 6-month milestone in formant frequency change is nuanced by an earlier shift observed specifically in positive vocalizations. This may reflect a more precocious expansion in the use of vowel-like sounds to express positive emotions, potentially facilitating early vocal play and exploration in a dyadic context. Such developments are particularly relevant during this stage of infancy, which is typically marked by dyadic interactions with caregivers - so-called ‘protoconversations’ - in which infants increasingly and actively adapt their vocal parameters to engage in reciprocal vocal exchanges with adult partners (Gratier & Devouche, 2011).

Emotional valence also influenced developmental changes in vocal quality and variability, albeit in a less straightforward manner. For negative and neutral vocalizations, a clear reduction in vocal variability features—such as jitter, shimmer, and spectral flux—was observed from 0–3 to 3–6 months. This aligns with research on the progressive refinement of vocal motor control (Nathani et al., 2006), and may also suggest a progressively more direct definition towards the acoustical properties of their native speech. However, this pattern stabilized into the 6–9 months range, with again a trend inversion for negative vocalizations after 9 months. Interestingly, these changes in timbre variability were not significant for positive vocalizations, which tended to be consistently instable across development. Echoing remarks about formant structure and variability, positive emotions being more likely to be reflected in vocal play during dyadic interactions, they may retain a consistent amount of instability even through later development stages. By 9–12 months, neutral vocalizations exhibit lower vocal instability - both in frequency and amplitude domains - as reflected in reduced jitter and shimmer values. Compared to both positive and negative vocalizations, this suggests that neutral sounds are marked by greater timbral stability and periodicity at this stage, whereas emotionally expressive vocalizations continue to display higher acoustic variability. This aligns with the idea that emotionally salient vocal production (both positive and negative) tend to preserve variability (Scherer et al., 2003), possibly as a means of communicative differentiation and efficacy. Overall, these findings may suggest that the developmental trajectory of vocal control—reflected in the balance between noise and periodicity captured by jitter and shimmer values—differs by emotional valence, supporting the idea that variability itself may serve an adaptive and functional role in affective communication early in life (Gerosa et al., 2007; Wermke et al., 2002). These interpretations should however be taken with caution considering the relatively messy patterns of these last acoustic parameters compared to the other PCA components.

As a probable consequence of these valence-specific developmental changes, vocalizations exhibited more precise emotional acoustic profiles in older vocalizations. The linear discriminant analysis showed most accuracy for vocalizations in the 6-9- and 9-12-months range, often correctly predicting both age and valence in these windows. As such, even considering the limits of experimenter categorization (see section below), infant prelinguistic vocalizing develops to be more than just shapeless sounds with context-dependent interpretations. Rather, they carry acoustic signatures reflecting the infants’ affective states that both underpin and are shaped by infants’ first vocal interactions.

Overall, these findings suggest that the developmental trajectory of infant vocalisations is shaped not only by age-related maturation but also by emotional valence, with vocalisations becoming progressively marked by valence-specific acoustic profiles rather than reflecting an emotionally undifferentiated stage.

### Limitations and future directions

The present study, while providing valuable insight into infant emotional communication, suffers from a few limitations. One such limit lays in the initial categorization of emotional valence, which ultimately depended on the appreciation of the experimenters; even the use of external cues such as caregiver responses implies a full and continuous mutual understanding between baby and caregivers, which is not guaranteed. Although this caveat is inevitable in in infant studies, future research should focus on methods to refine and consolidate the categorization of the present vocalization database, including perceptual validations by large panels of listeners. In addition, this research would benefit from an even larger and more diverse set of vocalizations, including analyses for each month of development, more specific emotion labels, and multi-cultural perspectives.

## Conclusion

This study provides evidence that the acoustic parameters of infant vocalizations encode emotional valence in emerging and consistent patterns, indicating that affective information is at least partially embedded in stable acoustic features from early infancy. These findings challenge the view of complete emotional neutrality in preverbal non-cry vocalizations and suggest that such vocalizations may function as early affective signals, offering caregivers meaningful acoustic cues alongside contextual information. The differentiation of vocalizations by valence enhances our understanding of how functional flexibility emerges through the developmental tuning of prosodic features. Furthermore, the results highlight the need for future research to determine whether adults can reliably infer emotional valence from acoustic features alone, and to what extent such cues serve as stable and complementary inputs in the interpretation of infant emotions. Such investigations are critical for elucidating the perceptual foundations of early socio-emotional communication and, from a broader perspective, for understanding how emotional encoding in the voice may evolve into more complex expressive forms, such as music and dynamic artistic performance.

