## Supplementary Materials for "Acoustic Features of Emotional Expression in Preverbal Infant Vocalizations"

**Table S1.** Results of the Principal Components Analysis on the GeMAPS acoustic parameters. Each row corresponds to an acoustic parameter as extracted by the openSMILE software. Each column is a rotated component from the PCA. For readability, only component loadings greater than 0.6 are displayed. The last three rows correspond to the overall loading scores and variance explained by each component. See Eyben et al. (2016) for a full description of each parameter.

| Parameter | RC1 | RC5 | RC2 | RC3 | RC4 |
| --- | --- | --- | --- | --- | --- |
| F0semitoneFrom27.5Hz_sma3nz_amean | 0.78 |  |  |  |  |
| F0semitoneFrom27.5Hz_sma3nz_percentile20.0 | 0.64 |  |  |  |  |
| F0semitoneFrom27.5Hz_sma3nz_percentile50.0 | 0.73 |  |  |  |  |
| F0semitoneFrom27.5Hz_sma3nz_percentile80.0 | 0.71 |  |  |  |  |
| mfcc1_sma3_amean | -0.84 |  |  |  |  |
| mfcc2_sma3_amean | -0.61 |  |  |  |  |
| logRelF0.H1.A3_sma3nz_amean | -0.66 |  |  |  |  |
| F1bandwidth_sma3nz_amean | -0.70 |  |  |  |  |
| F1bandwidth_sma3nz_stddevNorm | 0.61 |  |  |  |  |
| alphaRatioV_sma3nz_amean | 0.89 |  |  |  |  |
| hammarbergIndexV_sma3nz_amean | -0.78 |  |  |  |  |
| slopeV500.1500_sma3nz_amean | 0.79 |  |  |  |  |
| mfcc1V_sma3nz_amean | -0.85 |  |  |  |  |
| mfcc2V_sma3nz_amean | -0.61 |  |  |  |  |
| alphaRatioUV_sma3nz_amean |  |  |  |  |  |
| F0semitoneFrom27.5Hz_sma3nz_stddevNorm |  | 0.67 |  |  |  |
| F0semitoneFrom27.5Hz_sma3nz_pctlrange0.2 |  | 0.62 |  |  |  |
| spectralFlux_sma3_amean |  | 0.70 |  |  |  |
| jitterLocal_sma3nz_amean |  | 0.74 |  |  |  |
| shimmerLocaldB_sma3nz_amean |  | 0.74 |  |  |  |
| HNRdBACF_sma3nz_amean |  | -0.83 |  |  |  |
| spectralFluxV_sma3nz_amean |  | 0.71 |  |  |  |
| spectralFluxUV_sma3nz_amean |  |  |  |  |  |
| loudnessPeaksPerSec |  |  |  |  |  |
| VoicedSegmentsPerSec |  | 0.65 |  |  |  |
| loudness_sma3_amean |  |  |  |  |  |
| loudness_sma3_stddevNorm |  |  | 0.81 |  |  |
| loudness_sma3_percentile20.0 |  |  | -0.76 |  |  |
| loudness_sma3_percentile50.0 |  |  | -0.63 |  |  |
| spectralFlux_sma3_stddevNorm |  |  | 0.75 |  |  |
| F1amplitudeLogRelF0_sma3nz_amean |  |  | -0.75 |  |  |
| F2amplitudeLogRelF0_sma3nz_amean |  |  | -0.75 |  |  |
| F3amplitudeLogRelF0_sma3nz_amean |  |  | -0.74 |  |  |
| spectralFluxV_sma3nz_stddevNorm |  |  |  |  |  |
| MeanVoicedSegmentLengthSec |  |  |  |  |  |
| MeanUnvoicedSegmentLength |  |  | 0.61 |  |  |
| StddevUnvoicedSegmentLength |  |  | 0.61 |  |  |
| loudness_sma3_percentile80.0 |  |  |  |  |  |
| F1frequency_sma3nz_amean |  |  |  | -0.77 |  |
| F1frequency_sma3nz_stddevNorm |  |  |  | 0.64 |  |
| F2frequency_sma3nz_amean |  |  |  | -0.65 |  |
| F2frequency_sma3nz_stddevNorm |  |  |  | 0.72 |  |
| F3frequency_sma3nz_amean |  |  |  | -0.74 |  |
| F3frequency_sma3nz_stddevNorm |  |  |  |  |  |
| slopeV0.500_sma3nz_amean |  |  |  |  |  |
| mfcc4_sma3_amean |  |  |  |  | -0.61 |
| F2bandwidth_sma3nz_amean |  |  |  |  | -0.69 |
| F2bandwidth_sma3nz_stddevNorm |  |  |  |  | 0.67 |
| F3bandwidth_sma3nz_amean |  |  |  |  | -0.67 |
| F3bandwidth_sma3nz_stddevNorm |  |  |  |  |  |
| mfcc4V_sma3nz_amean |  |  |  |  | -0.64 |
| F0semitoneFrom27.5Hz_sma3nz_meanRisingSlope | | | | | |
| F0semitoneFrom27.5Hz_sma3nz_stddevRisingSlope | | | | | |
| F0semitoneFrom27.5Hz_sma3nz_meanFallingSlope | | | | | |
| F0semitoneFrom27.5Hz_sma3nz_stddevFallingSlope | | | | | |
| loudness_sma3_pctlrange0.2 |  |  |  |  |  |
| loudness_sma3_meanRisingSlope |  |  |  |  |  |
| loudness_sma3_stddevRisingSlope |  |  |  |  |  |
| loudness_sma3_meanFallingSlope |  |  |  |  |  |
| loudness_sma3_stddevFallingSlope |  |  |  |  |  |
| mfcc1_sma3_stddevNorm |  |  |  |  |  |
| mfcc2_sma3_stddevNorm |  |  |  |  |  |
| mfcc3_sma3_amean |  |  |  |  |  |
| mfcc3_sma3_stddevNorm |  |  |  |  |  |
| mfcc4_sma3_stddevNorm |  |  |  |  |  |
| jitterLocal_sma3nz_stddevNorm |  |  |  |  |  |
| shimmerLocaldB_sma3nz_stddevNorm |  |  |  |  |  |
| HNRdBACF_sma3nz_stddevNorm |  |  |  |  |  |
| logRelF0.H1.H2_sma3nz_amean |  |  |  |  |  |
| logRelF0.H1.H2_sma3nz_stddevNorm |  |  |  |  |  |
| logRelF0.H1.A3_sma3nz_stddevNorm |  |  |  |  |  |
| F1amplitudeLogRelF0_sma3nz_stddevNorm |  |  |  |  |  |
| F2amplitudeLogRelF0_sma3nz_stddevNorm |  |  |  |  |  |
| F3amplitudeLogRelF0_sma3nz_stddevNorm |  |  |  |  |  |
| alphaRatioV_sma3nz_stddevNorm |  |  |  |  |  |
| hammarbergIndexV_sma3nz_stddevNorm |  |  |  |  |  |
| slopeV0.500_sma3nz_stddevNorm |  |  |  |  |  |
| slopeV500.1500_sma3nz_stddevNorm |  |  |  |  |  |
| mfcc1V_sma3nz_stddevNorm |  |  |  |  |  |
| mfcc2V_sma3nz_stddevNorm |  |  |  |  |  |
| mfcc3V_sma3nz_amean |  |  |  |  |  |
| mfcc3V_sma3nz_stddevNorm |  |  |  |  |  |
| mfcc4V_sma3nz_stddevNorm |  |  |  |  |  |
| hammarbergIndexUV_sma3nz_amean |  |  |  |  |  |
| slopeUV0.500_sma3nz_amean |  |  |  |  |  |
| slopeUV500.1500_sma3nz_amean |  |  |  |  |  |
| StddevVoicedSegmentLengthSec |  |  |  |  |  |
| equivalentSoundLevel_dBp |  |  |  |  |  |
| **SS loadings** | 11.06 | 8.25 | 7.29 | 6.18 | 5.31 |
| **Proportion Variance** | 0.13 | 0.09 | 0.08 | 0.07 | 0.06 |
| **Cumulative Variance** | 0.13 | 0.22 | 0.30 | 0.37 | 0.43 |

**Table S2.** Mean and standard deviation for the component scores of each RC, for each Age Range and Valence condition.

| Age | Valence | RC1 M (SD) | RC2 M (SD) | RC3 M (SD) | RC4 M (SD) | RC5 M (SD) |
| --- | --- | --- | --- | --- | --- | --- |
| 0-3 | Negative | .23(.66) | .33(.92) | -.37(.69) | .46(.58) | .51(.98) |
| 0-3 | Neutral | -.48(.71) | -.21(.98) | -.25(.82) | .64(.67) | .42(.92) |
| 0-3 | Positive | -.47(.77) | -.57(.77) | -.02(.73) | .69(.58) | .09(.75) |
| 3-6 | Negative | -.22(1.05) | .40(.52) | -.77(.91) | .36(.95) | -.23(.83) |
| 3-6 | Neutral | -.46(.57) | .36(.65) | -.72(1.15) | .07(.99) | -.46(1.00) |
| 3-6 | Positive | -.34(.95) | -.08(1.03) | .11(.84) | .26(.78) | .09(1.00) |
| 6-9 | Negative | .37(1.25) | .29(1.02) | .83(.81) | .11(.82) | -.27(.96) |
| 6-9 | Neutral | -.22(1.13) | -.17(1.20) | .36(.99) | .01(1.05) | -.11(1.38) |
| 6-9 | Positive | .16(1.12) | -.53(1.11) | .70(.78) | -.26(.95) | -.11(.69) |
| 9-12 | Negative | .76(.77) | .39(.98) | .25(1.04) | -.62(.93) | .26(1.00) |
| 9-12 | Neutral | .05(.72) | .02(1.09) | -.19(1.02) | -.78(1.02) | -.39(.79) |
| 9-12 | Positive | .60(1.12) | -.23(.97) | .08(.82) | -.94(.92) | .21(1.14) |
